# Utilizing top-down hyperspectral imaging for monitoring genotype and growth conditions in maize

**DOI:** 10.1101/2020.01.21.914069

**Authors:** Sara B. Tirado, Susan St Dennis, Tara A. Enders, Nathan M. Springer

## Abstract

There is significant enthusiasm about the potential for hyperspectral imaging to document variation among plant species, genotypes or growing conditions. However, in many cases the application of hyperspectral imaging is performed in highly controlled situations that focus on a flat portion of a leaf or side-views of plants that would be difficult to obtain in field settings. We were interested in assessing the potential for applying hyperspectral imaging to document variation in genotypes or abiotic stresses in a fashion that could be implemented in field settings. Specifically, we focused on collecting top-down hyperspectral images of maize seedlings similar to a view that would be collected in a typical maize field. A top-down image of a maize seedling includes a view into the funnel-like whorl at the center of the plant with several leaves radiating outwards. There is substantial variability in the reflectance profile of different portions of this plant. To deal with the variability in reflectance profiles that arises from this morphology we implemented a method that divides the longest leaf into 10 segments from the center to the leaf tip. We show that using these segments provides improved ability to discriminate different genotypes or abiotic stress conditions (heat, cold or salinity stress) for maize seedlings. We also found substantial differences in the ability to successfully classify abiotic stress conditions among different inbred genotypes of maize. This provides an approach that can be implemented to help classify genotype and environmental variation for maize seedlings that could be implemented in field settings.

**Significance Statement:** This study describes the importance of using spatial information for the analysis of hyperspectral images of maize seedling. The segmentation of maize seedling leaves provides improved resolution for using hyperspectral variation to document genotypic and environmental variation in maize.

## Introduction

Abiotic stresses cause major yield declines across many crops and can limit production by up to 70% (Boyer, 1982). Advances in molecular tools have greatly facilitated breeders in efficiently identifying and selecting germplasm with favorable traits such as tolerance to abiotic stresses; however, breeders still rely on obtaining high quality phenotypic data for developing and implementing these methods (Masuka et al., 2012). Phenotyping has become the main bottleneck in making breeding advances because current methods of phenotyping involve a large amount of time and labor. This limits their applications across breeding programs which typically consist of large populations comprised of thousands of lines grown in replicates across multiple environments (Myles et al., 2009). To effectively breed for tolerance to abiotic stresses, quantifying the severity of the response to a particular stress across different genotypes as well as their ability to recover from the stress is crucial. This would require temporal measurements of phenotypes linked to the stress response which increases the complexity in making progress in breeding for such traits.

The development of high throughput phenotyping tools has taken surge over the last couple of years to obtain phenotype data quickly and at low costs. Most of these methods rely on remote sensing techniques that utilize sensors to capture images of plants and subsequently processing the images to extract meaningful traits. Sensors that measure different ranges of the electromagnetic spectrum have been applied in agriculture. RGB imaging has been widely used to extract morphological traits linked to plant productivity across different crop species in field and indoor settings (Watanebe et al., 2017; Feng et al., 2018; Varela et al., 2017; Enders et al., 2019). Thermal imaging and near infrared combined with visible imaging have been used to extract information of drought stress across grasses and legume crops (Martynenko et al., 2016; Biju et al., 2018; Benavente et al., 2013; Jin et al., 2017; Zhang et al., 2012).

More recently, with the advent of advanced machine and deep learning algorithms, multispectral and hyperspectral sensors that generate large amounts of data at very high spectral and spatial resolutions have been applied in four key areas in plant phenotyping: identification, classification, quantification, and prediction of a particular stress (Singh et al., 2015). With hyperspectral imaging, the user can take advantage of hundreds of spectral channels to uncover materials and biochemical processes, such as the degradation of pigment molecules and changes in water content, within plant tissues that can differentiate and potentially quantify differences across species, genotypes, and stresses. The degradation of pigments such as chlorophyll alters the amount of reflected, absorbed, and transmitted radiation and can therefore be passively captured using spectral imaging (Blackburn, 2007).

Hyperspectral imaging has been applied for the identification and quantification of several bacterial and fungal infections including fusarium head blight and leaf rust in wheat (Alisaac et al., 2019; Mahlein et al., 2019; Qiu et al., 2019; Ashourloo et al., 2014; Bauriegel et al., 2011), powdery mildew in barley (Thomas et al., 2018), *Cercospora* leaf spot, powdery mildew and leaf rust in sugarbeet (Mahlein et al., 2012; Rumpf et al., 2010) and *Sclerotinia sclerotiorum* in oilseed rape plants (Kong et al., 2018). Diseases caused by bacterial or fungal infections tend to have characteristic features such as bacterial pustules with neighboring chlorotic tissue or necrotic lesions that are picked up and easily distinguished using spectral imaging. However, identifying more subtle symptoms such as those caused by abiotic stresses can be challenging. Pandey et al. (2017) found hyperspectral imaging to be useful in quantifying plant leaf chemical properties that could aid in detecting water and nutrient deficiencies among crops and Obeidat et al. (2018) discovered that spectral indices correlated with chlorophyll content could help distinguish between genotypes and cold-stressed plants in indoor settings. Similarly, Behmann et al. (2014) found that hyperspectral imaging can be used to cluster barley plant pixels into different levels of drought-stress based on amount of chlorosis and senescence. Also, Römer et al. (2012) was able to detect drought stress early in development for cereal grains in both indoor and field settings based on a matrix factorization technique that allows for the computation of how similar a plant pixel is to the typical spectrum of a healthy plant. Another way to identify subtle abiotic stress signals using spectral imaging is by correlating reflectance data with other more laborious, time-consuming or costly measurements correlated to the stress response. Feng et al (2019) made a link between hyperspectral measurements of okra leaves with measurements linked to leaf chlorophyll content and fresh weight traditionally used to assess salt stress across crops.

A common problem when analyzing spectral data of plant surfaces is taking into account uneven light scattering that occurs upon the interaction between incident light and the plant surface being captured (Makdessi et al., 2017). Plant material possesses non-Lambertian reflectance properties and plants themselves contain a large amount of morphological variation causing differences in angle relative to the sensor across leaf segments. Many studies that have evaluated the use of hyperspectral imaging for assessing plant abiotic stresses have utilized indoor setups with uniform, nonreflective backgrounds and have dismissed the effects of plant morphology by securing the plant leaves on a flat background (Obeidat et al., 2018); however, this limits their application in natural settings. Other studies have proposed ways to account for plant architectural variation. Behman et al. (2015) proposed a method to account for differential light scattering by performing geometric calibration of hyperspectral cameras that connects a 3D model with a 2D image and Mohd Asaari et al. (2018) applied a standard normal variate normalization method to correct spectra for uneven illumination effects. Moghimi et al. (2018) circumvented plant architectural differences by identifying endmembers indicative of all plant pixels for a given line in a given treatment and used these to identify salt stress across wheat lines. Feng et al. (2019), on the other hand, was able to develop an instance segmentation model using deep learning to segment individual okra plant leaves for further evaluation, which is a suitable approach for crops where leaves lie relatively flat horizontally with respect to the sensor.

Variation in plant architecture across different crops species can make finding a single approach for analyzing spectra data challenging; however, it can also be taken advantage of in the context of finding discriminatory patterns within an individual crop. Upon accounting for differences in light scattering of different plant surfaces, the large number of plant pixels representing single or multiple individuals are commonly reduced to a single value such as an endmember (Moghimi et al., 2018) or an average (Mohd Asaari et al., 2018). Looking at all the plant pixels throughout the plant can elucidate biochemical processes in response to certain stimuli that vary spatially throughout a plant. This spatial variation could be useful for identifying more subtle symptoms that may be masked out by reducing the data to a single value per line or treatment or by normalizing the data to account for scattered light due to differences in plant morphology. Moreover, although multiple studies have identified indices that are useful for a particular stress, they have not looked into how these would change due to plant morphology. This study aims to elucidate the effects of morphology and stress on the spatial variation of reflectance values within plant leaves and compare the ability of reflectance data for different regions of the leaf to resolve genetic and environmental factors relative to reflectance data for the entire plant. Currently, this remains unknown and could enlighten new mechanisms for identifying, classifying, quantifying and predicting the onset and recovery of biotic and abiotic stresses where little variation is observed with the naked eye.

## Results and discussion

In most field settings, hyperspectral images of cereal crops are collected from above, resulting in a top-down view of the plants. We sought to develop approaches for the analysis of hyperspectral images for maize plants that could be applicable to field settings. We obtained raw intensity data using a top-down approach for wavelengths ranging from approximately 400 to 1000 nm for several controlled-condition experiments. The experiments contained maize seedlings of multiple inbred genotypes subjected to different environmental treatments. Plants were illuminated by halogen lights, which are oriented in two parallel rows of bulbs on either side of the camera and each image contained 3 plants (Figure S1). Raw intensity values for the resulting images of plants were converted to reflectance using white and dark references and then normalized by their L2 norm (see methods). For each plant, the NDVI values were utilized to identify pixels containing plant tissue and thresholded to generate a binary image mask to extract the reflectance values at each wavelength for entire plants.

This approach was applied to several different experiments that are summarized in Table S1. We sought to address different themes in our analyses of this data. First, different genotypes or environments often result in changes in plant morphology (Enders et al., 2019). We evaluated the potential of hyperspectral data to capture morphological differences by utilizing changes in reflectance for specific regions of a plant. In our analyses, we compared the ability of using whole plant data relative to using specific regions of leaves in the ability to resolve genetic or environmental factors. Second, hyperspectral imaging provides opportunities to identify genetic or environmental variation. We assessed the relative ability of using hyperspectral imaging data to accurately classify environmental conditions in different genotypes using machine-learning approaches. To achieve these goals, we collected hyperspectral data using the described system for two experiments. The first experiment (E1) consists of three replicates of five genotypes of maize seedlings under four treatment groups (control, cold, salt, and heat) imaged at a single time point. The chosen genotypes were previously demonstrated to differ in responses to cold stress (Enders et al.,2019). The second experiment (E2) consists of two genotypes under four treatment groups (control and three severities of salt stress) imaged at three timepoints (immediately before the stress, 2 days following the stress treatment, and 4 days after following the stress treatment). A total of 540 plant images are represented in the dataset. We have used the dataset to address a series of questions about the ability of hyperspectral imaging to resolve the effects of growth stage, genotype and various abiotic stresses across portions of maize leaves.

### Spatial variation in top-down images of seedlings

The system and approach that was used to generate images of maize seedlings results in a relatively large number of pixels (1,908-13,831) for each plant. These pixels exhibit a range of reflectance values with substantial standard deviation (Figure S2). A top-down image of a maize seedling consists of a central whorl from which leaves extend. The whorl has a funnel-like shape with each leaf extending in an arc (Figure 1A). Given the variation in reflectance based on the orientation of the plant surface relative to the lights and camera, there is substantial variability in reflectance from the central whorl to the leaf blade tip. In addition, there is biological variation in gene expression and physiological properties of leaves from the base to the tip (Li et al., 2010). We sought to compare the average normalized reflectance values from each plant within zones extracted along the length of a leaf. To classify relatively consistent zones of a plant leaf, we implemented an approach to divide the longest leaf into ten sections and identify the plant pixels within each section (Figure 1, see methods for details). This resulted in a set of 10 segments that were used as a mask for the hyperspectral image cube to extract normalized reflectance values by leaf zone.

**Figure 1.**
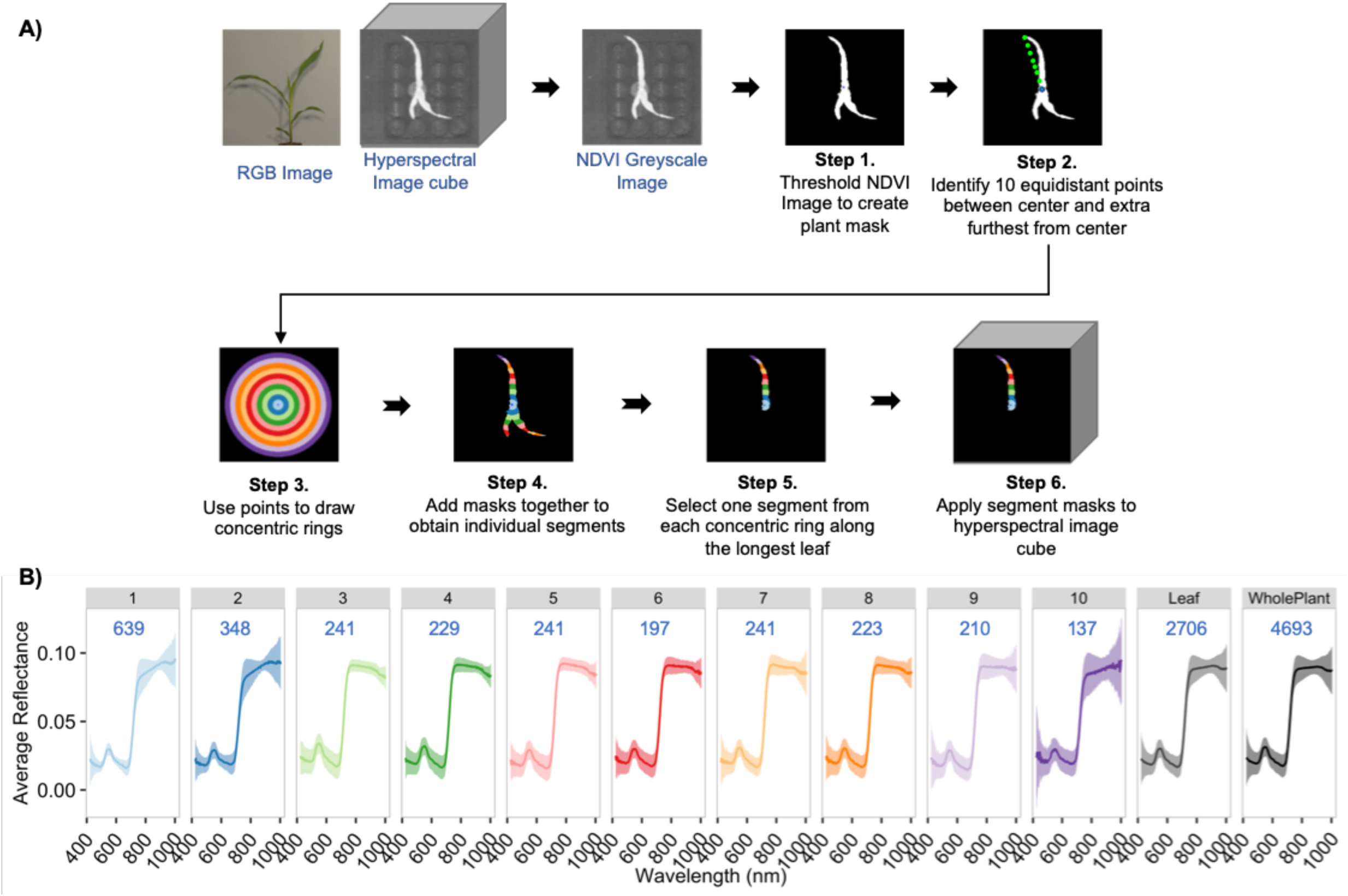
Evaluating individual leaf segments. **A)** Procedure for extracting hyperspectral data from ten segments across the longest leaf of each plant. **B)** Mean (dark line) and variance in reflectance of individual leaf segments, the entire longest leaf (all leaf segments combined) and the entire plant of a Mo17 control plant at 13 days after sowing. Pixel counts represented in each panel are indicated by the blue text. Leaf segment is indicated at the top of each panel in black text.

The average reflectance profiles and variance were compared for the entire leaf relative to each of the ten leaf sections. Substantial variability is observed in pixel counts across leaf segments due to differences in width across the length of the leaf (Figure 1). While the overall pattern of reflectance values is generally similar among plant segments there is substantial variability within and across plant segments for the magnitude of specific patterns. In many cases the variation in spectral profiles across leaf segments can be difficult to visualize when using the reflectance patterns for the entire spectrum (Figure 1B). PCA of the reflectance values for each segment of each plant suggests differences in the most outer segments of the leaves relative to more central regions of the leaf (Figure 2A). A comparison of the reflectance profiles across leaf segments focused only on the visible range of the spectrum reveals distinct reflectance profiles between leaf segments near the center or leaf tip relative to the middle portion of the leaf in both Mo17 and PH207 (Figure 2B).

**Figure 2.**
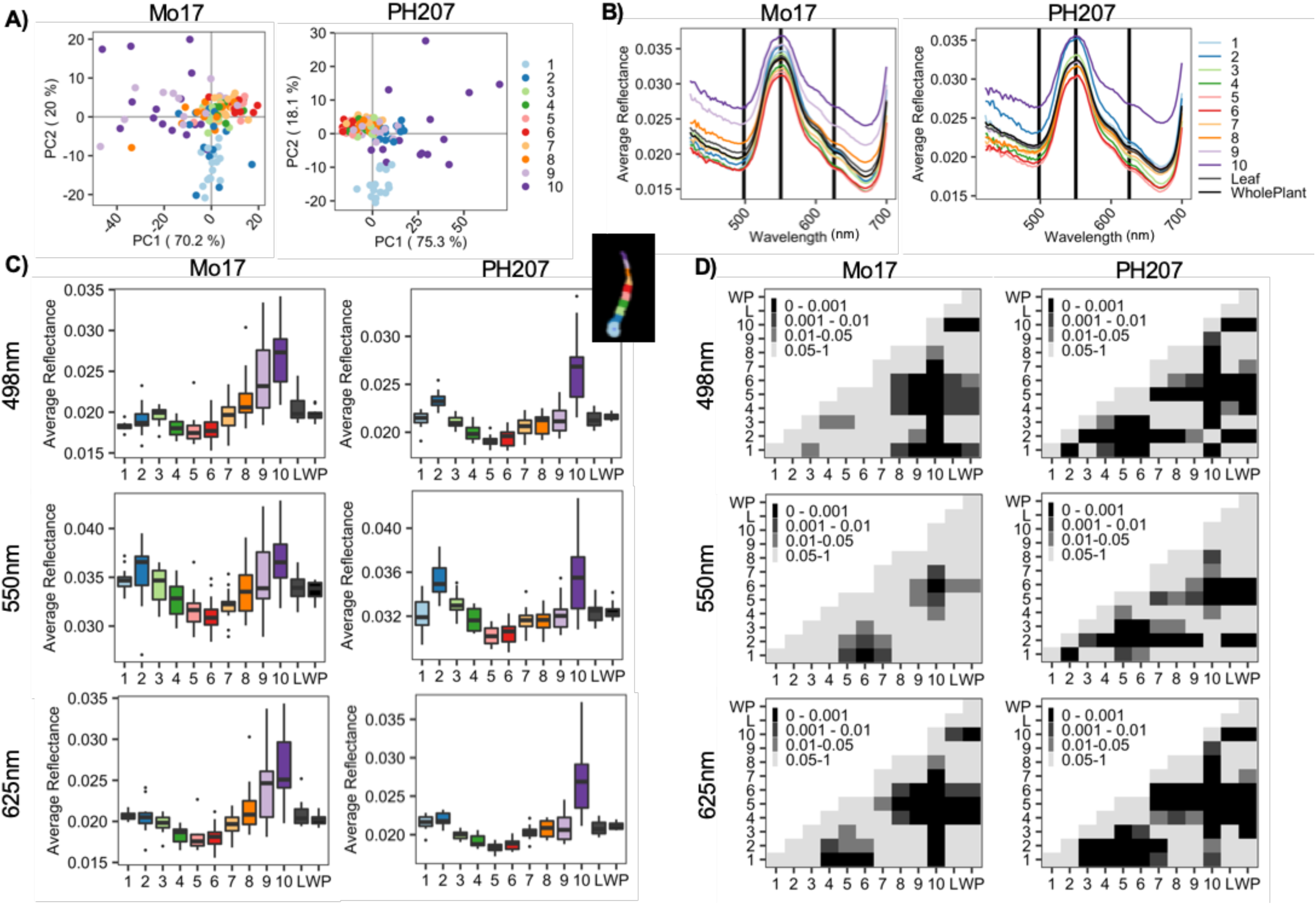
Reflectance of leaf segments across all Mo17 and PH207 control plants in experiment E2 at 15 days after sowing. **A)** PCA biplots of the mean reflectance across all wavelengths of all 10 leaf segments for individual plants. Each datapoint constitutes an individual leaf segment for a single plant. **B)** Mean reflectance for the 10 leaf segments, the entire leaf and the entire plant. All plants were utilized to calculate the mean. Vertical black lines indicate wavelengths at 498 nm, 550 nm, and 625 nm. **C)** Distributions of the average reflectance per plant of leaf segments, the entire leaf, and the entire plant at 498 nm, 550 nm and 625 nm. **D)** Adjusted p-values from a pairwise wilcox test between pairwise comparisons of leaf segment, entire leaf (L), and whole plant (WP) reflectance, at 498 nm, 550 nm and 625 nm. Results were adjusted for multiple comparisons using the “holm” method. Black color indicates p-value < 0.001.

We selected three representative wavelengths in the red, green, and blue range of the light spectrum (625 nm, 550 nm, and 498 nm; Figure 2B) and assessed the distribution of average reflectance values for all plants for each of the leaf segments as well as the entire leaf (Figure 2C). For several of the segments the reflectance values for these three wavelengths exhibit distributions of values that are significantly different from each other or from the entire leaf (Figure 2D). In general, the patterns are similar for the two genotypes. Relative to the values observed for the entire leaf there are often significant differences in the distribution of values seen from the middle and the tip of the leaf. The tip of the leaf is often distinct from many other zones as well. These observations highlight the variability throughout a single leaf and suggest that using all values for a plant or leaf will likely obscure spatial variation that may occur due to developmental, genetic or environmental factors.

### Stable patterns of hyperspectral signal for different stages of seedling growth

The differences in hyperspectral profiles were assessed for PH207 seedlings grown in control conditions that were 11, 13 and 15 days after sowing to document whether there are differences as seedlings mature and whether the differences among leaf segments are consistent over time (Figure 3). PCA reveals that differences across leaf segments account for most of the observable variation in reflectance intensity compared to differences observed between days (Figure 3A). Across the three time points, average reflectance values cluster into groups corresponding to leaf segments near the leaf tip, leaf segments towards the middle of the leaf, and leaf segments towards the center whorl. No clustering by date was observed even though plant size and morphology changes were observed based on trait data obtained from RGB images (Figure S3, see methods). The profiles of reflectance values are slightly different on the three dates (Figure 3B) but the distribution patterns for the different leaf segments relative to each other remain consistent. Examination of the distribution of values at three wavelengths reveals similar distributions for plants at the three dates and similar trends among the different leaf segments (Figure 3C).

**Figure 3.**
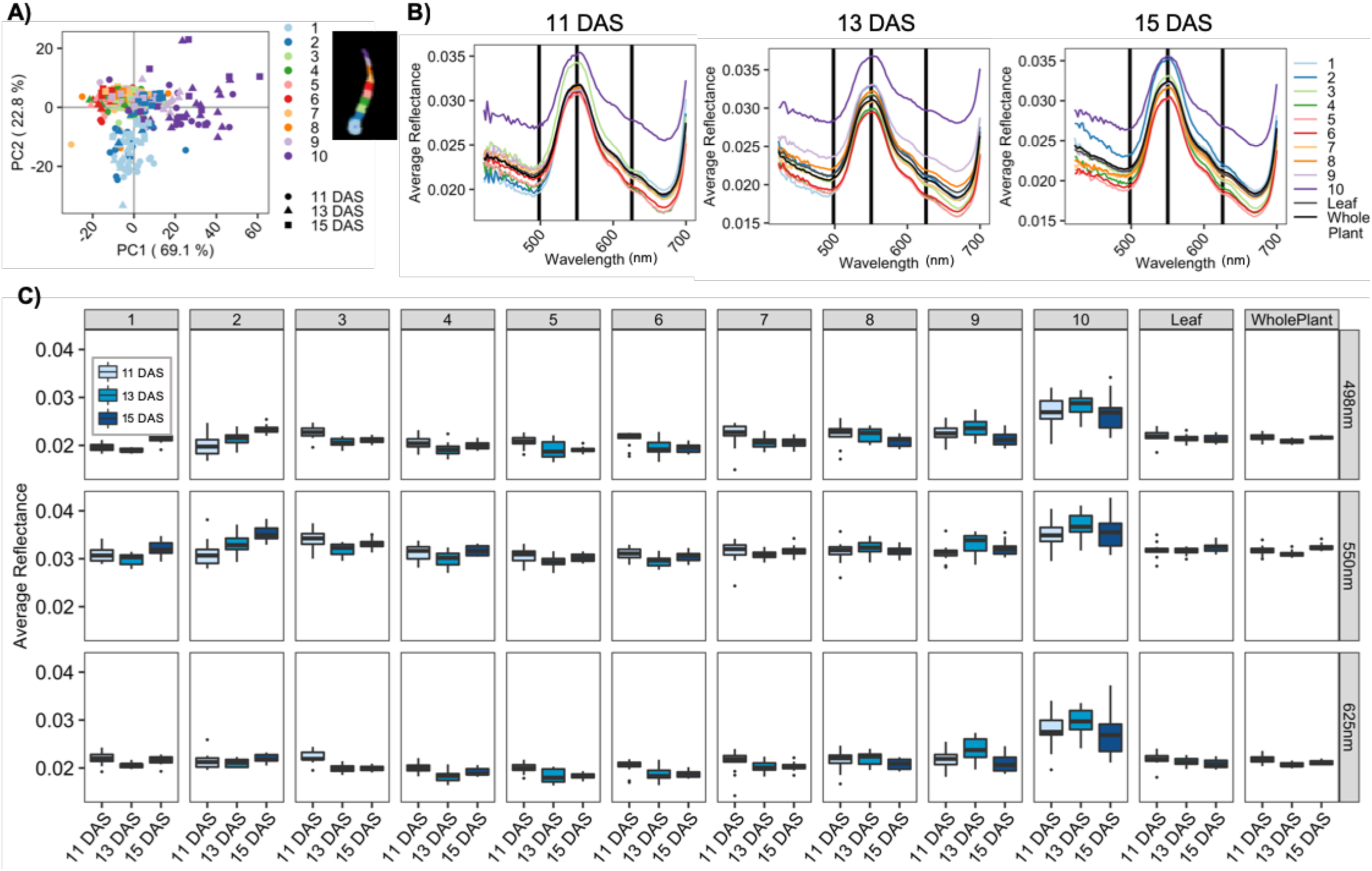
Reflectance of leaf segments across all PH207 control plants in experiment E2 across 11, 13 and 15 days after sowing. **A)** PCA biplots of the mean reflectance across all wavelengths of all 10 leaf segments for individual plants. Each datapoint constitutes an individual leaf segment for a single plant. **B)** Mean reflectance for the 10 leaf segments, the entire leaf and the entire plant. All plants were utilized to calculate the mean. Vertical black lines indicate wavelengths at 498 nm, 550 nm, and 625 nm. **C)** Distributions of the average reflectance per plant of leaf segments, the entire leaf, and the entire plant at 498 nm, 550 nm and 625 nm across the three days of imaging.

### Ability to distinguish genotypes using hyperspectral imaging

Hyperspectral profiling has been widely used for separating different plant species such as weeds (Pantazi, Moshou and Bravo, 2016) and tropical forest trees (Laybros et al., 2019). Fewer studies have used hyperspectral profiling to separate different genotypes or lines of the same species. A study by Obeidat et al. (2018) showed that genotype main effects across short-season maize lines significantly contributed to variation in various spectral reflectance indices as well as spectral reflectance in the visible and near-infrared range when comparing hyperspectral scans of flat leaves. This variation, particularly in the spectral reflectance across the visible range of the spectrum, was likely due to chlorophyll and carotenoid differences (Obeidat et al., 2018). To assess variation in spectral reflectance among maize genotypes, we applied our leaf segmentation approach to compare the reflectance values across individual leaf segments among the different inbred lines grown in control conditions in experiment E1. Experiment E1 consisted of images from 9 plants for five different genotypes. While the overall average profiles for the entire leaf are relatively similar in the visible range, there are some leaf segments that show more pronounced differences among genotypes (Figure 4A). In particular, MS71 shows higher reflectance for wavelengths near 550 nm for central segments of the leaf but lower reflectance compared to other genotypes for wavelengths near 500 nm and 675 nm for segments near the leaf tip. However, these differences are reduced in averages that include all pixels for the entire leaf (Figure 4A).

**Figure 4.**
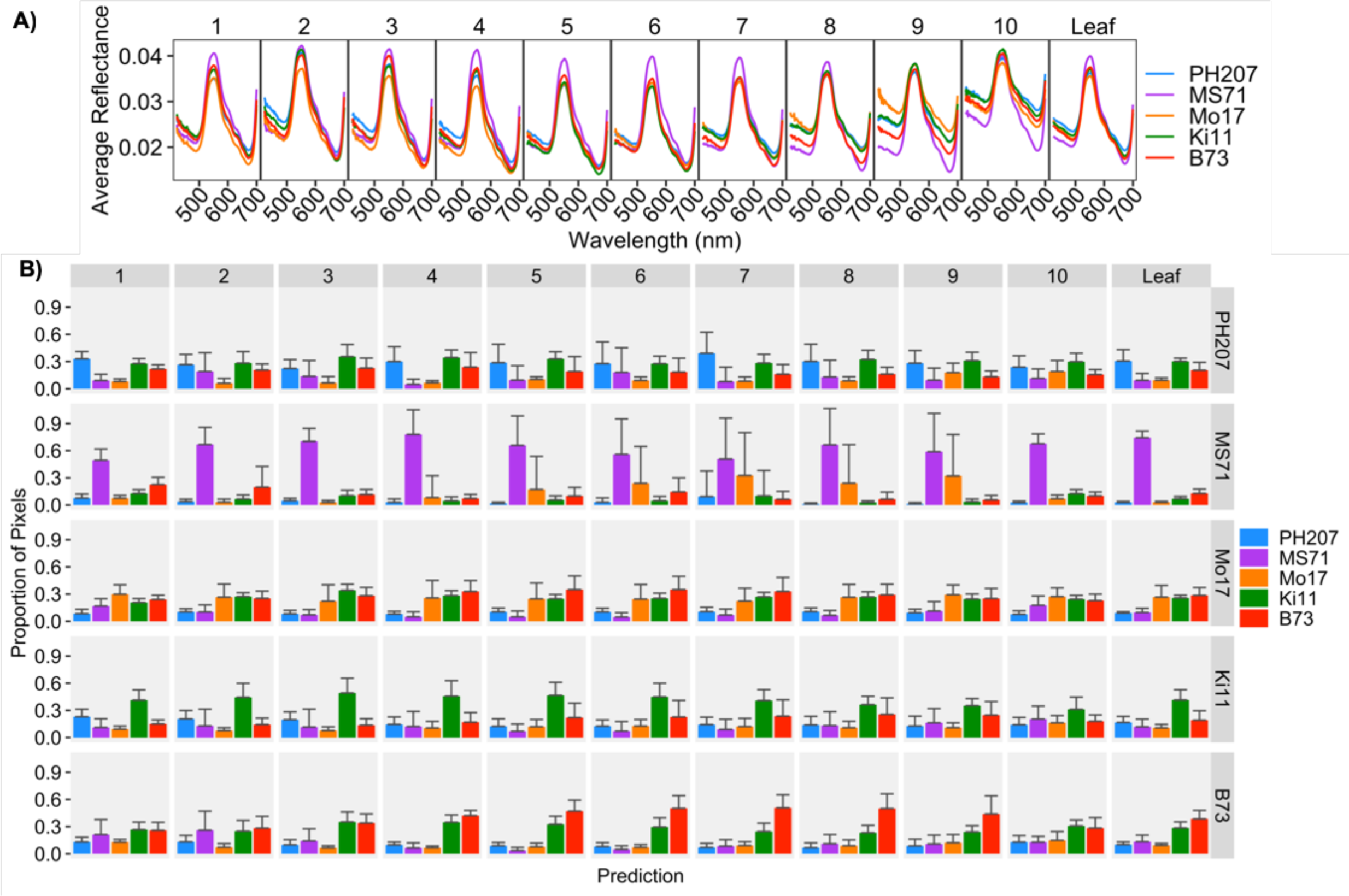
Genotypic differences in hyperspectral profiles for all control plants of each genotype in Experiment E1. **A)** Average reflectance values across all control plants of each genotype. **D)** Average proportion of pixels per plant from all plants of a certain genotype (rows) for each leaf segment (columns) predicted to belong to a certain genotypic class. Error bars indicate the standard deviation in pixel values from the mean.

To further evaluate whether specific leaf segments can be more useful than whole leaves in distinguishing individual genotypes, we developed a cubic support vector machine (SVM) model utilizing □ of all the pixel values from all plants grown in control conditions in experiment E1 to predict the corresponding genotype. The predictor variables included normalized reflectance values for wavelengths in the visible range as well as the leaf segment of the corresponding pixel. We then applied this model to predict the genotype of the remaining □ pixel values in the dataset. This included pixels from all leaf segments. The proportion of pixels per plant that were classified into each genotype for each of the zones of each maize line was determined (Figure 4B). The model is able to correctly identify MS71 pixels in most leaf segments although the accuracy is somewhat lower in the whorl region (Figure 4B). The other genotypes are less accurately identified. B73 is most accurately identified for the central leaf segments but is often confused with Ki11 (Figure 4B). PH207 and Mo17 generally exhibit relatively low correct prediction accuracies throughout the leaf and are frequently mis-classified as B73 or Ki11 (Figure 4B).

### Ability to distinguish and quantify abiotic stresses using hyperspectral imaging

Two different experiments were performed to investigate the potential to utilize hyperspectral profiling for documenting the effects of abiotic stress on maize seedlings. For experiment E2 we treated two genotypes, Mo17 and PH207, with three different concentrations of salt applied on day 11 immediately after imaging (Figure 5A). These plants were then imaged two and four days after the stress application. Two cubic SVM models were developed using □ of all pixels from the control and the medium salt stress treatment groups. In both models, the treatment (Control or 0.75M NaCl) was the response variable. The first model was trained using pixels randomly selected throughout the entire plant; however, the second model only contained pixels randomly selected from the longest leaf. In the first model, the normalized reflectance values for wavelengths in the visible range, the genotype, and the day of imaging represented as days after sowing (DAS) were set as the predictor variables. In the second model, the corresponding leaf segment was also included as a predictor variable. Moreover, the first model was used to predict all the pixels from entire plants in the E2 dataset from all treatment groups into belonging to the control or salt stressed class (Figure 5B). On the other hand, the second model was used to predict all pixels from the longest leaf of all plants (Figure 5C).

**Figure 5.**
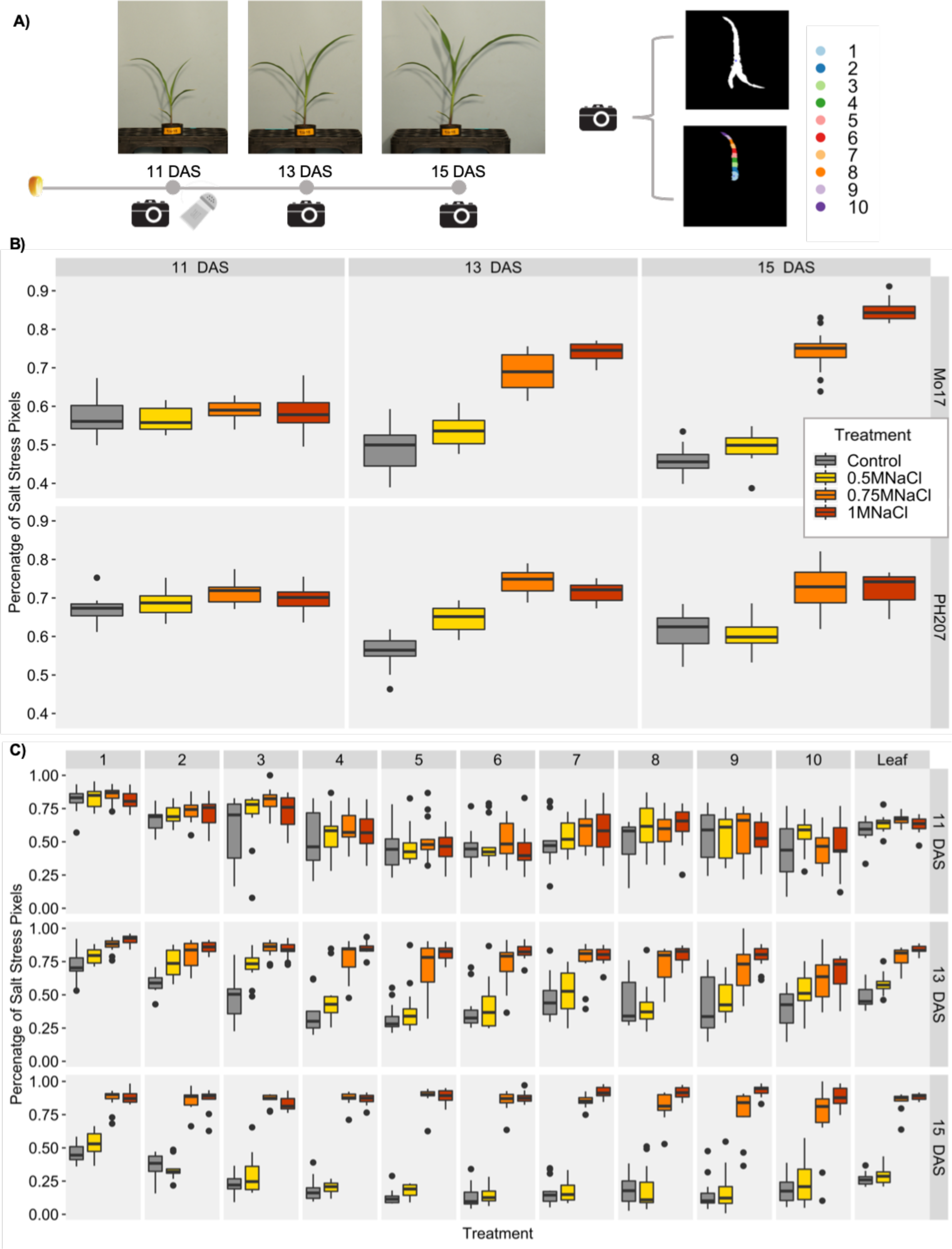
Proportion of pixels per plant classified as being salt stressed based on cubic SVM model developed from the wavelengths in the visible range of the spectrum and genotype as the predictor variables and using the medium salt stress and control treatments as the response variables. **A)** Three salt treatments were applied after imaging at 11 days after sowing (DAS). Plants were then imaged at 13 and 15 DAS. Reflectance values were calculated for entire plants, entire leaves and individual leaf segments. **B)** Proportion of total pixels for each plant across treatments in Experiment E2 classified as belonging to the salt stress category for the model developed using pixels across the whole plant. **C)** Proportion of total pixels for individual leaf segments and the entire leaf per plant for Mo17 classified as belonging to the salt stress category for a model developed using pixels across individual plant segments with segment number as a predictor variable.

As expected, the proportion of pixels classified as salt-stressed was not different for the control and the treatment groups at day 11 for either model as these images were collected prior to the actual stress treatment application (Figure 5B; Figure 5C). However, at day 13 and day 15, two and four days after application of salt we see substantial increases in the proportion of pixels classified as salt stressed (Figure 5B; Figure 5C). When looking at the classifications based on the entire plant using the first model, the proportion of pixels classified as salt stressed increases at higher concentrations of salt treatment and is higher at day 15 than at day 13 (Figure 5B). However, there are a high frequency of pixels in control plants classified as salt stressed in this analysis. A comparison of the predictive ability for different leaf segments revealed substantial variation across the leaf. At days 13 and 15 the segments from the middle of the leaf have higher correct prediction accuracy than segments near the leaf tip or whorl (Figure 5C). Importantly, these mid-leaf segments also outperform the predictions based on using the entire plant. A relatively small proportion of pixels from control plants are classified as stressed in these mid-leaf segments while the majority of pixels in plants with 0.75M or 1M NaCl treatment are classified as stressed.

Experiment E1 included five genotypes treated with four conditions including control, heat stress, cold stress and salt stress and plants were imaged after two days of the stress treatment. Visual examination of the plants revealed differences in severity of stress response for the different genotypes (Figure S4). This is quantified for cold stress by Enders et al (2019). For example, Ki11 tends to have strong responses, especially to cold and salt stress while Mo17 has minimal visual responses to the stresses (Figure S4). The average hyperspectral profiles for the entire leaf reveal limited changes for Mo17 but some differences for Ki11 (Figure 6A). The differences in hyperspectral profiles for the different treatments were more severe in some leaf segments compared to others (Figure 6A). A cubic SVM model was developed to predict the treatment using predictor variables of normalized reflectance values, genotype, and leaf segment. The proportion of pixels classified into each condition is shown for each segment of each actual treatment (Figure 6B, C). For Ki11 there is a high true prediction accuracy for cold and salt stress across all segments of the plants; however, the prediction accuracy is further improved for segments near the middle of the leaf for cold stress and the tip of the leaf for salt stress (Figure 6B). Heat stress is not predicted as accurately with substantial confusion between control and heat stress (Figure 6B). This likely reflects minimal phenotypic response to heat stress for Ki11. Similar patterns of enhanced prediction accuracy utilizing middle segments of the leaf compared to the entire leaf across treatments are also observed for the other four genotypes (Figure S5, S6).

**Figure 6.**
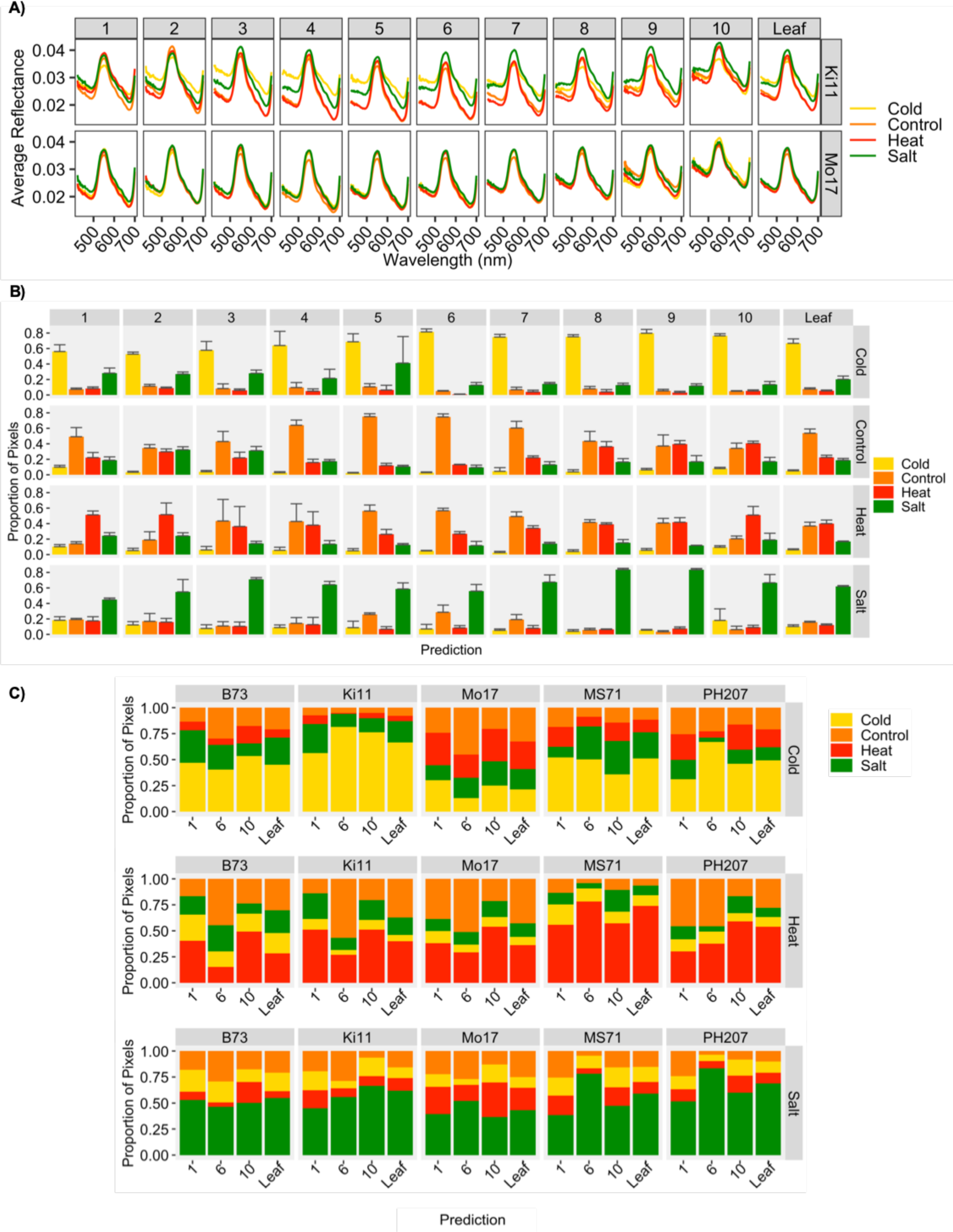
Differences in hyperspectral profiles across treatments for all Ki11 and Mo17 plants in Experiment E1. **A)** Average reflectance values for all plants in the cold, control, heat and salt stress treatments of Ki11 and Mo17 in Experiment E1. **B)** Average proportion of pixels from all Ki11 plants of a certain treatment (rows) for each leaf segment (columns) predicted to belong to a certain treatment class. Bars represent the mean proportion of all plants per category and error bars represent the standard deviation around the mean. **C)** Average proportion of pixels from all plants of a certain treatment (rows) for genotypes across selected leaf segments predicted to belong to a certain treatment class. Bars represent the mean proportion of all plants per genotype and treatment class predicted to belong to a certain treatment.

If we compare the accuracies of a representative leaf segment in the center whorl, the middle portion of the leaf, the tip of the leaf, and the entire leaf in predicting the abiotic stress response across genotypes, we observe differences in utility of different leaf segments based on the stress being predicted as well as the genotype (Figure 6C). Leaf segments in the middle portion of the leaf as well as the leaf tip provided the highest true prediction accuracy overall across treatments and genotypes compared to the center whorl and utilizing all segments form the leaf. Leaf segments towards the middle portion of the leaf (segment 6 in this case) provide a higher prediction accuracy across most genotypes for predicting cold stress and across some genotypes (Mo17, MS71 and PH207) for salt stress; however, the leaf tip was more informative for heat stress and across some genotypes such as Ki11 for salt stress (Figure 6C). Overall, most pixels that were misclassified across genotypes and treatments were predicted to belong to the control treatment group.

## Conclusions

Hyperspectral profiling provides new opportunities for optical analysis of trait variation in crops. Many studies have reported the ability to monitor physiological changes in plants using point profiles of reflectance from a single region of a leaf (Meacham-Hensold et al., 2019; Silva-Perez et al., 2018; Smith et al., 2004; Yendrek et al., 2017). However, there has been less analysis of the ability to separate effects of genotype or environmental conditions using whole plant images. In this study we highlight the potential for using hyperspectral imaging but also show that using averages of whole plants provides less resolution than focusing on specific regions of plants. The variation in spectral profiles from the base of the leaf to the tip of the leaf likely represents a combination of physiological differences as well as variation in the plant shape/leaf angle resulting in differing reflectance. In this study, we have not separated these factors but instead have simply relied upon segmentation of the leaf to reduce variance and improve discrimination.

There are several limitations to our approach for segmenting the longest leaf and making comparisons of specific leaf segments across dates, genotypes and treatments. In this work, we performed manual detection of the center of the plant and the remainder of the leaf identification and segmentation process was automated. It is likely that spectral properties or plant shape could be used for automated detection of the center of the whorl. Additionally, when comparing leaf segments for varying genotypes or growing conditions the length of the longest leaf may vary as plants exhibit different growth rates for different stress conditions or among genotypes. This results in differing numbers of pixels for the segments being compared. However, since we are segmenting into 10 equally sized regions the relative segmentation of the leaf should remain consistent. Another potential issue is the variation in the angle of the leaf tip. As a leaf emerges from the whorl it has an upright angle. As the leaf extends the tip will shift from upright to horizontal to having a downward angle. There is biological variation among plants at the same developmental stage for the angle of the leaf tip. This may result in increased variance for segments near the leaf tip, as noted in our PCA plots (Figure 2A, 3A). However, many stress conditions have visible effects on the leaf margins near the leaf tip and this region provided the best classification for some stresses. One additional potential complication is the presence of mixed pixels that include some plant tissue as well as background. We implemented relatively strict cutoffs to minimize the number of mixed pixels obtained after plant segmentation but there are likely a small number of mixed pixels captured in our plant masks. These may be represented in uneven quantities across leaf segments with more mixed pixels appearing in narrower segments such as the leaf tip relative to the base of leaf.

Our findings highlight the utility of plant segmentation for improved accuracy of genotype or environment predictions using hyperspectral data. It is worthwhile to note that there is not a single region of the leaf with the highest performance. Instead the most accurate regions varied for different stresses or genotypes. The use of wider panels of genotypes would likely result in classification of groups with similar behaviors, but in this study we focused on improving the methods for stress detection in a small set of variable genotypes. Our classification prediction rates vary substantially. This is likely due to variation among the genotypes. Some genotypes are more tolerant of certain abiotic stresses and we can observe a higher proportion of pixels misclassified into the control classes for these. In contrast, genotypes that are more sensitive to a certain stress exhibit a larger prediction accuracy for stress prediction for the particular treatment. The approaches of segmenting leaves for hyperspectral analysis can likely be conducted for larger scale field experiments and may improve the utility of hyperspectral profiles for documenting genotype, environment and genotype by environment effects.

## Methods

### Plant growth

Two experiments were conducted, E1 and E2 (Table S1). For all experiments, seeds were planted approximately 2 inches below the surface in 40 cubic inch D□40 DeePots (Stuewe and Sons, Inc.) containing a 1:1 mix of SunGro (Agawam, MA) horticulture professional growing mix and autoclaved field soil. All plants were grown in Conviron growth chambers with a 16 hr 30°C and 8 hr 20°C day/night cycle and watered every other day.

In experiment E1, five maize genotypes (B73, Mo17, PH207, Ki11, MS71) were subjected to four treatment conditions (control, cold, heat, salt). Plants for all treatments were grown in control conditions until 11 DAS when the stresses were applied for the cold, heat and salt treatments. The cold□stress treatments were implemented using a Thermo Scientific refrigerated incubator programmed with a 16 hr 6°C and 8 hr 2°C day/night cycle and applied for 48 hours. The heat□stress treatments were implemented using a Thermo Scientific refrigerated incubator programmed with a 16 hr 39°C and 8 hr 29°C day/night cycle and applied for 48 hours. The salt stress was a single 50 mL 0.75 M NaCl treatment at Zeitgeber Time 2 (ZT2) at 11 DAS. Plants were imaged with our hyperspectral and RGB imaging systems at the end of the stress treatment at ZT2 at 13 DAS. Three experimental replicates were grown each consisting of three plants per genotype per treatment condition, for a total of 9 plants per genotype per treatment.

Experiment E2 consisted of two maize genotypes (Mo17 and PH207) subjected to four treatment conditions (control, low salt, medium salt and high salt stress). The low, medium and high salt stress treatments were implemented by a single 50 mL 0.5 M NaCl, 0.75 M NaCl or 1 M NaCl treatment, respectively, at ZT2 at 11 DAS. Plants were imaged with our hyperspectral and RGB imaging systems before undergoing stress at ZT2 at 11 DAS, at the end of the stress treatment at ZT2 at 13 DAS and two days after the stress treatment at 15 DAS. The experiment consisted of 15 plants per genotype per treatment.

### Hyperspectral image acquisition

To capture the hyperspectral images, a custom-built line-scanning system from Middleton Spectral Vision (Madison, WI) was utilized (Figure S1). The system contains a Specim V10 spectrograph with a spectral range of 400 to 1000 nm and approximately a 1 nm spectral resolution. The spectrograph was mounted on an Imperx IPX-2M30 camera with 1600×1200 pixel spatial resolution. When acquiring hyperspectral images, a spectral binning of 2x was applied to obtain an average spectral resolution of 2.3nm.

### Hyperspectral Data Pre-processing and Normalization

Black and white references are gathered by capturing and averaging 10 hyperspectral frames with the camera shutter closed and for a white lambertian reference panel each date of data collection. Intensity values for each pixel at each wavelength are converted to reflectance values by subtracting the dark reference and dividing the result by the difference between the white and dark references (Yoon and Park, 2015). The resulting reflectance values are then normalized by dividing each spectrum by its L_2 norm, or the square root of the sum of the squares of that signature, following the equation

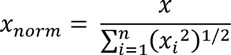

where *x* is the full vector of reflectance data in image, *i* is the response band, *n* is the total number of measured wavelengths and *x_i_* is the full vector of reflectance data in image for the response band *i*.

### Segmenting Plant Material and Longest Leaf into Individual Segments

All approaches for identifying, segmenting and extracting data from leaf segments were implemented utilizing custom MATLAB algorithms (MATLAB, 2018a). Plant material was segmented by calculating the NDVI value of each pixel and thresholding to a value of 0.35 or greater to create a binary plant segmentation mask. This threshold was found to balance maintaining the highest percentage of plant pixels while minimizing the number of mixed pixels in the extracted data. Each hyperspectral image contained three plants of the same genotype and treatment in a defined location. Individual plant objects were identified from the plant material mask using the *bwconncomp* function in MATLAB which returns connected components with a connectivity of 8 (MATLAB, 2018b). Objects caused by background noise were then removed by only keeping objects that had a minimum of 1,000 pixels and a maximum of 3,000 pixels and allocated an ID (plant A, B or C) based on their location in the image.

For each plant (plant A, B and C) of each image, the approximate center of the leaf whorl was identified by manually identifying the x-y coordinates of the plant center from an NDVI grayscale representation. This represented the only manual input in the pipeline. Extrema of the plant object were then automatically identified using the *regionprops* function in MATLAB (MATLAB, 2018b). Using the x-y coordinates for each terminal extrema and whorl center, the extrema farthest away from the center was identified for each plant representing the tip of the longest leaf. The distance of the center to the longest leaf tip was then divided into ten equidistant points in linear space and 10 concentric rings were generated utilizing the identified distance between points as the radius (Figure 1A). Each ring segment was used as a mask coupled with the plant material segmentation mask to extract leaf segments along the plant (Figure 1A). To ensure that only segments belonging to a single, constant leaf were kept, only segments that also overlapped with a straight line that extended from the center of the plant to the longest leaf tip were kept. Each of these ten segments were used as a mask for the hyperspectral image cube, and reflectance data was extracted for wavelengths 420 nm to 1000 nm after trimming off noisy wavelengths. The reflectance values of each pixel in the plant were then normalized by the L2-norm calculated on a whole image basis for each wavelength (see Hyperspectral Data Pre-processing and Normalization methods section). This normalized data was then exported for further analysis.

### Outlier Detection and Removal

Individual leaf segments for each plant were visually assessed by looking at the leaf segment binary masks and all data from the given segment excluded in the analysis if the segment encompassed multiple leaves (which happened in cases where part of the given segment was close to the whorl before leaves separated or when leaves overlapped each other), if the leaf segment had less than 25 pixels, or if the segment was not on the primary longest leaf (which occurred in some cases where the leaf curled or overlapped another leaf). A total of 156 plants of the entire 540 had at least one leaf segment excluded (29%); however, the final number of leaf segments excluded was 187 out of the total 5400 (3%). The majority of excluded segments were located adjacent to the whorl in cases where the leaves were short and the second leaf segment encompassed multiple leaves.

### Prediction Model Development and Implementation

All cubic support vector machine (SVM) models were developed using the Classification Learner application in MATLAB with specified response and predictor variables (MATLAB, 2018c). Cubic SVM models were selected after testing 13 different machine learning algorithms as these models consistently provided the highest prediction accuracy across the different applications specified in this study. A random subsampling of 1/6th of all pixels from all plants in the target dataset was used as the training dataset to create the model (Figure S7). Five-fold cross-validation was utilized to evaluate the performance of the algorithm during model building to prevent model overfitting during training (Figure S7). This involved further randomly partitioning the data into a testing and training set five times. The training sets were used to train the supervised learning algorithm and the testing sets were used to obtain an average cross-validation error estimate to evaluate the algorithm performance. Each trained cubic SVM model was then exported and used to predict response classes for remaining 5/6th of pixels constituting the validation set and obtain prediction accuracy estimates (Figure S7).

### Statistical Analyses

The average spectra for each leaf segment of each plant was compared to the average spectra of each other segment as well as the average spectra for the entire leaf and the entire plant to assess which leaf segments significantly differ from each other. The comparisons were made by performing a pairwise Wilcoxon Rank Sum Test between all pairwise comparisons of leaf segments using the pairwise.wilcox.test function from the R stats package (R Core Team, 2012).

### RGB Trait Data Acquisition

RGB side-view images of each set of plants were collected immediately following hyperspectral data collection using the procedures specified in Enders et al. (2019).

The scripts and processes used to perform the data normalization and leaf segmentation are available at https://github.com/SBTirado/HS_LeafSegmentation.git.

## Acknowledgements

This work was supported by the National Science Foundation Plant Genome Award IOS-1444456.

## Data Statement

All hyperspectral datasets are available through CyVerse. Scripts utilized for processing the hyperspectral images as well as for leaf segmentation are available at https://github.com/SBTirado/HS_LeafSegmentation.git.

---

**Table S1.**
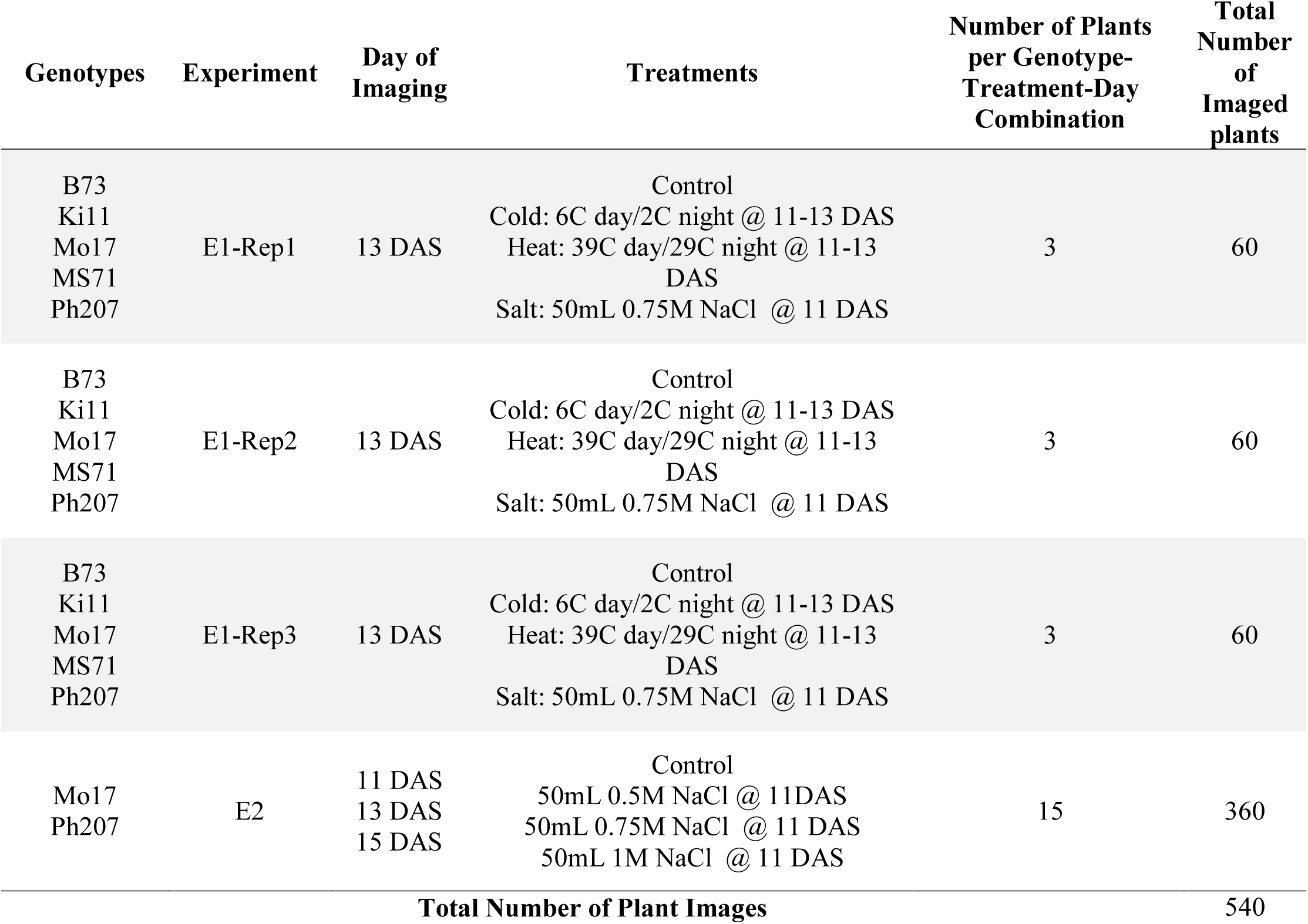
Summary of Experiments.

**Figure S1. Hyperspectral imaging setup. A)** Imaging system utilized. **B)** RGB representation of staged plants **C)** Sensor and light configuration.

**Figure S2. Variation in reflectance measurements among pixels within whole plants.** Standard deviation of pixel reflectance values across whole plants for three (A, B and C) Mo17 and PH207 control plants from a single plot imaged at the indicated day after sowing (DAS). Black line indicates reflectance at 625 nm.

**Figure S3. Mean RGB trait values for all PH207 control plants in experiment E2 across 11, 13 and 15 days after sowing (DAS).**

**Figure S4. RGB images for one representative plant in experiment E1 for each treatment for Ki11 and Mo17 genotypes 13 days after sowing.**

**Figure S5.** Average proportion of pixels from all Mo17 (**A**) and B73 (**B**) plants of a certain treatment (rows) for each leaf segment (columns) predicted to belong to a certain treatment class. Bars represent the mean proportion of all plants per category and error bars represent the standard deviation around the mean.

**Figure S6.** Average proportion of pixels from all MS71 (**A**) and PH207 (**B**) plants of a certain treatment (rows) for each leaf segment (columns) predicted to belong to a certain treatment class. Bars represent the mean proportion of all plants per category and error bars represent the standard deviation around the mean.

**Figure S7. SVM model training and testing procedure.**

