## Supplemental Figures for "Utilizing top-down hyperspectral imaging for monitoring genotype and growth conditions in maize"

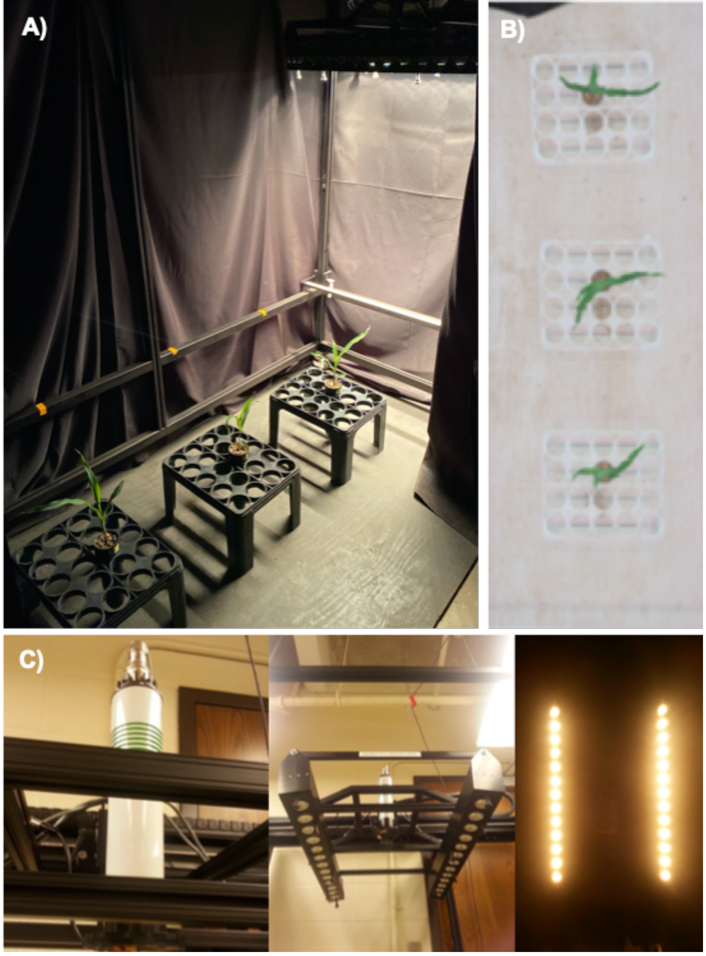


**Figure S1. Hyperspectral imaging setup. A)** Imaging system utilized. **B)** RGB representation of staged plants **C)** Sensor and light configuration.

**Table S1. Summary of experiments.**

| Genotypes | Experiment | Day of Imaging | Treatments | Number of Plants per Genotype-Treatment-Day Combination | Total Number of Imaged plants |
|---|---|---|---|---|---|
| B73<br>Ki11<br>Mo17<br>MS71<br>Ph207 | E1-Rep1 | 13 DAS | Control<br>Cold: 6C day/2C night @ 11-13 DAS<br>Heat: 39C day/29C night @ 11-13 DAS<br>Salt: 50mL 0.75M NaCl @ 11 DAS | 3 | 60 |
| B73<br>Ki11<br>Mo17<br>MS71<br>Ph207 | E1-Rep2 | 13 DAS | Control<br>Cold: 6C day/2C night @ 11-13 DAS<br>Heat: 39C day/29C night @ 11-13 DAS<br>Salt: 50mL 0.75M NaCl @ 11 DAS | 3 | 60 |
| B73<br>Ki11<br>Mo17<br>MS71<br>Ph207 | E1-Rep3 | 13 DAS | Control<br>Cold: 6C day/2C night @ 11-13 DAS<br>Heat: 39C day/29C night @ 11-13 DAS<br>Salt: 50mL 0.75M NaCl @ 11 DAS | 3 | 60 |
| Mo17<br>Ph207 | E2 | 11 DAS<br>13 DAS<br>15 DAS | Control<br>50mL 0.5M NaCl @ 11DAS<br>50mL 0.75M NaCl @ 11 DAS<br>50mL 1M NaCl @ 11 DAS | 15 | 360 |
| | | | | **Total Number of Plant Images** | 540 |

**A)**

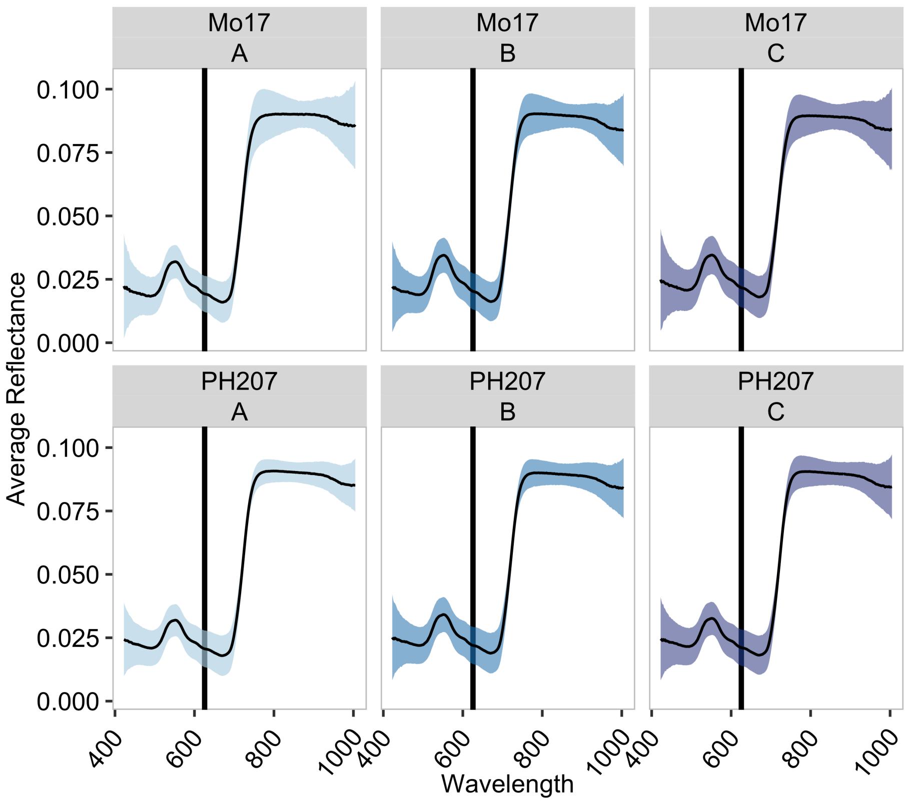


**Figure S2. Variation in reflectance measurements among pixels within whole plants.** Standard deviation of pixel reflectance values across whole plants for three (A, B and C) Mo17 and PH207 control plants from a single plot imaged at the indicated day after sowing (DAS). Black line indicates reflectance at 625 nm.

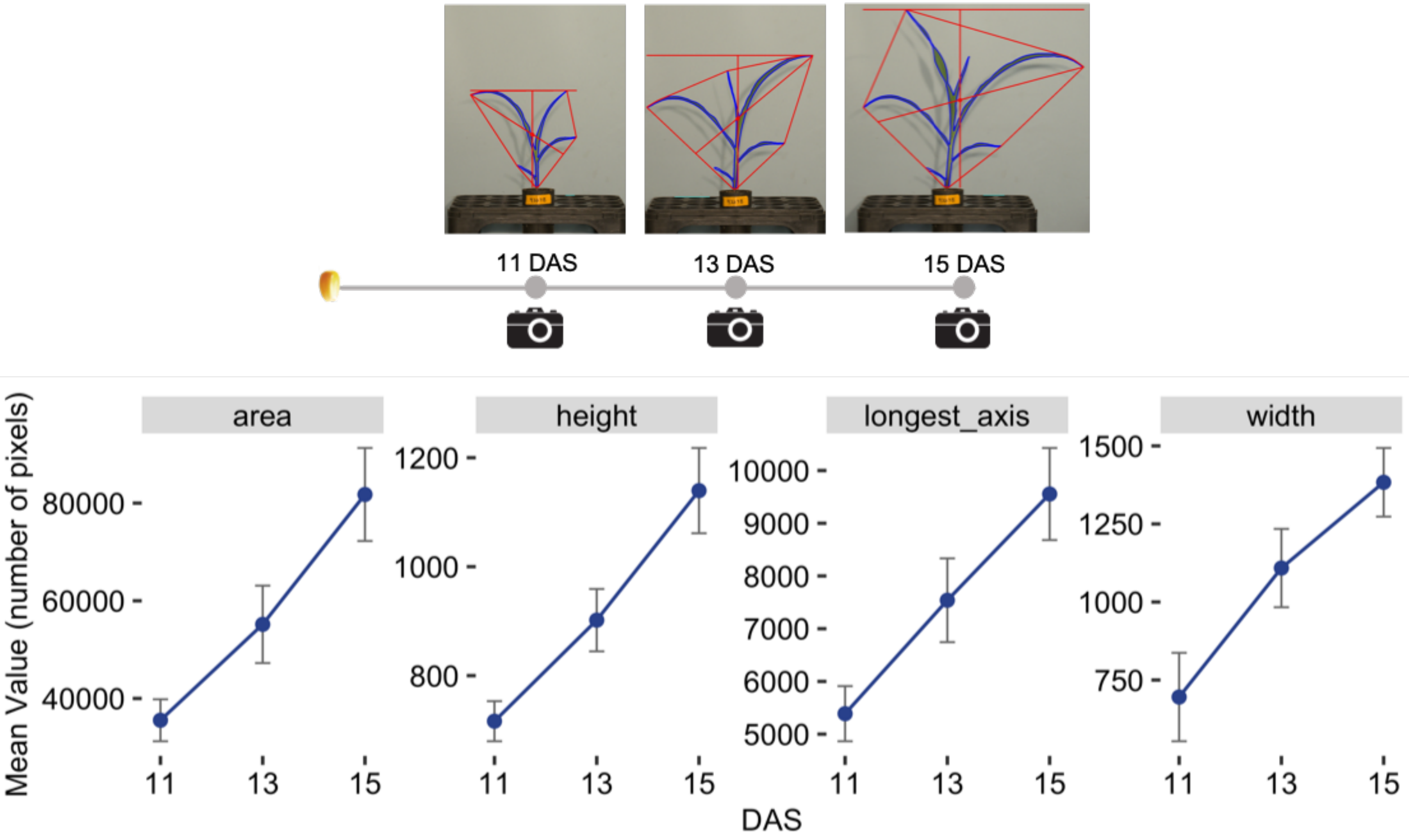


**Figure S3. Mean RGB trait values for all PH207 control plants in experiment E2 across 11, 13 and 15 days after sowing (DAS).**

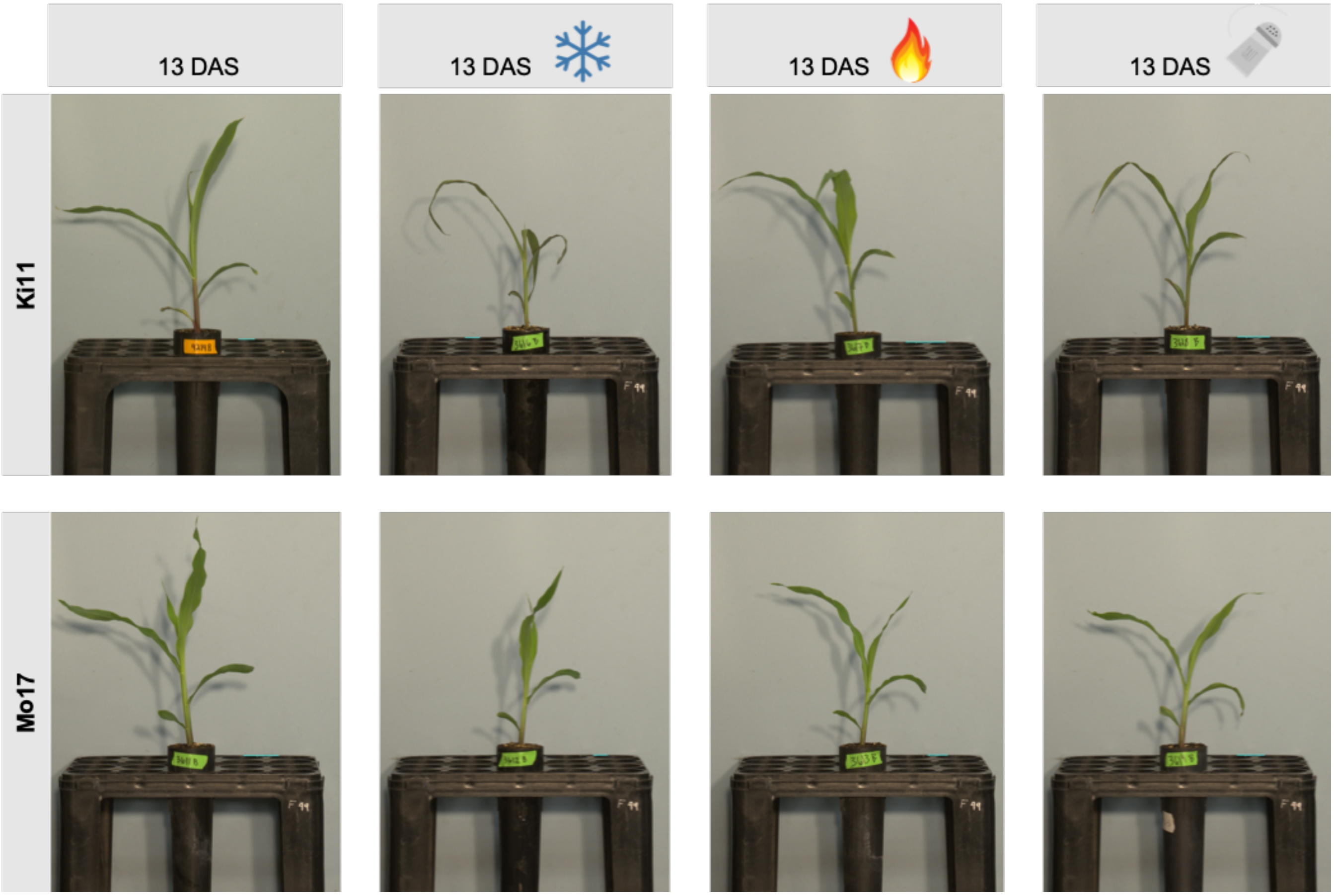


**Figure S4. RGB images for one representative plant in experiment E1 for each treatment for Ki11 and Mo17 genotypes 13 days after sowing.**

A)

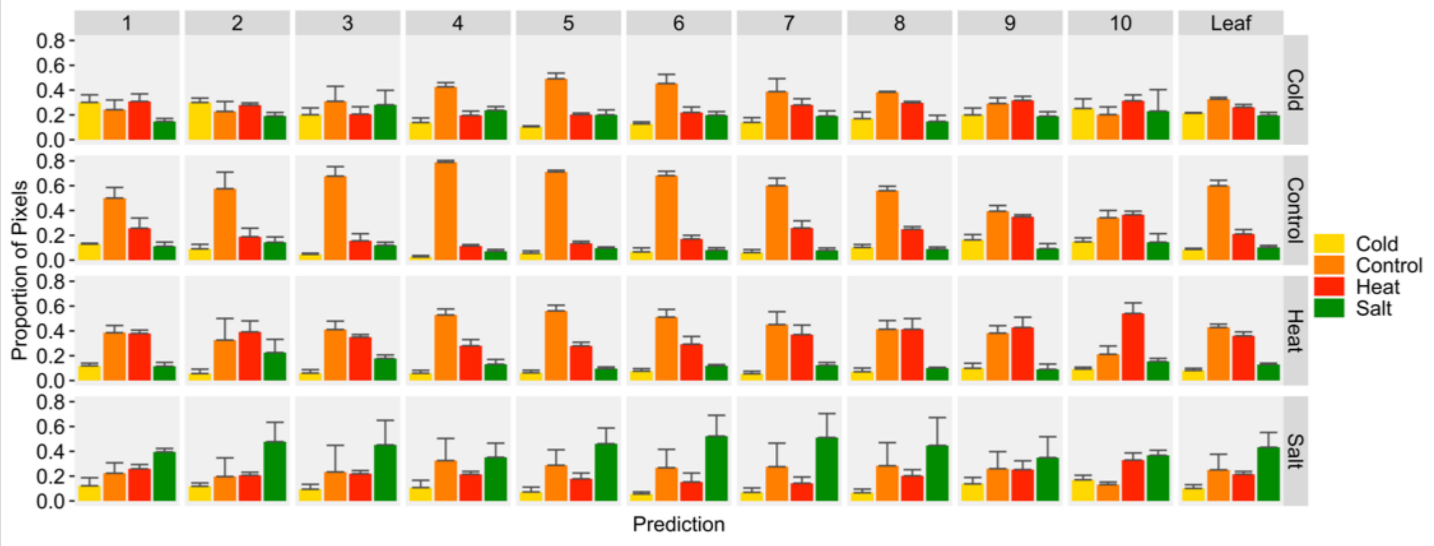


B)

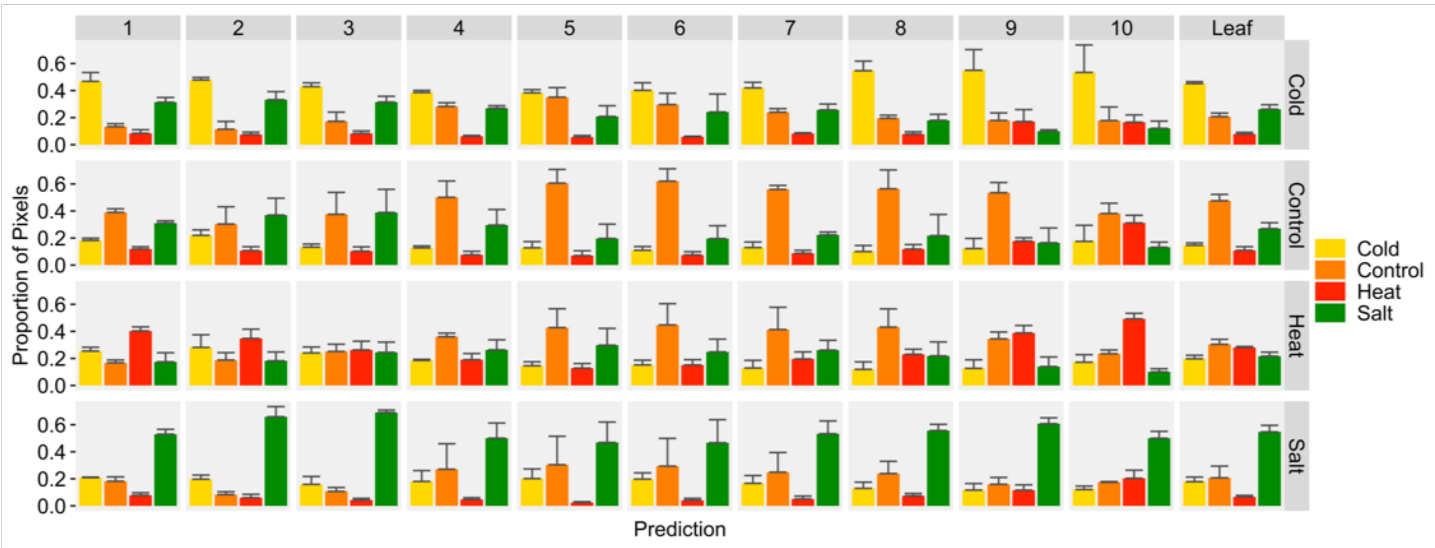


**Figure S5.** Average proportion of pixels from all Mo17 (**A**) and B73 (**B**) plants of a certain treatment (rows) for each leaf segment (columns) predicted to belong to a certain treatment class. Bars represent the mean proportion of all plants per category and error bars represent the standard deviation around the mean.

A)

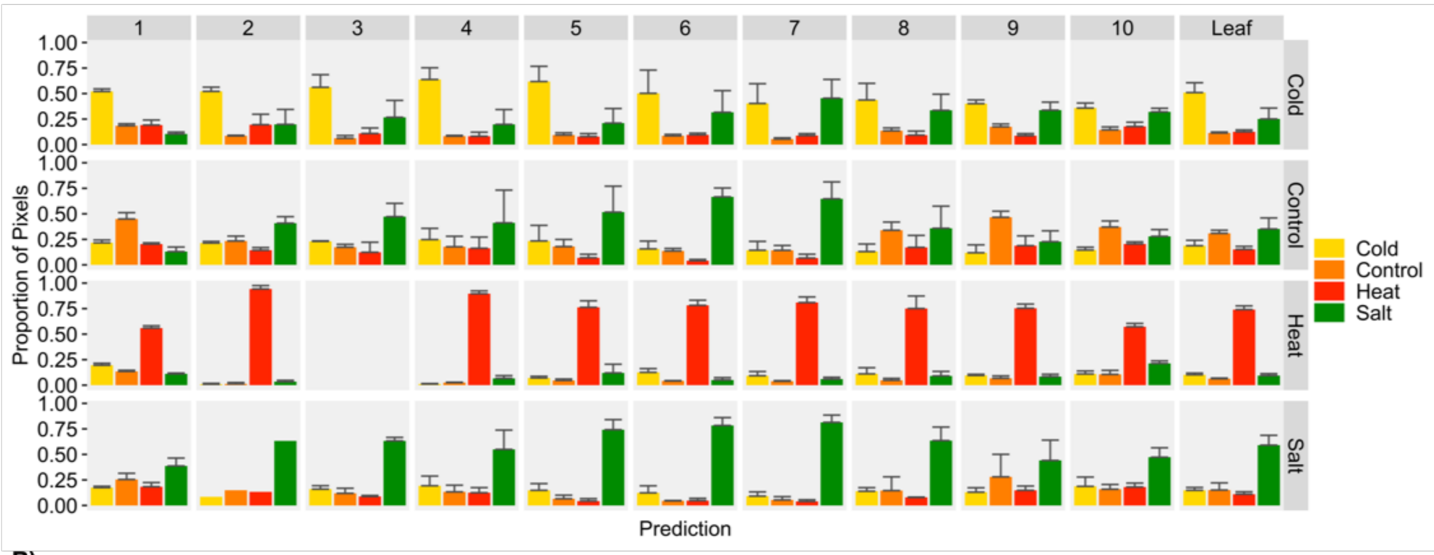


B)

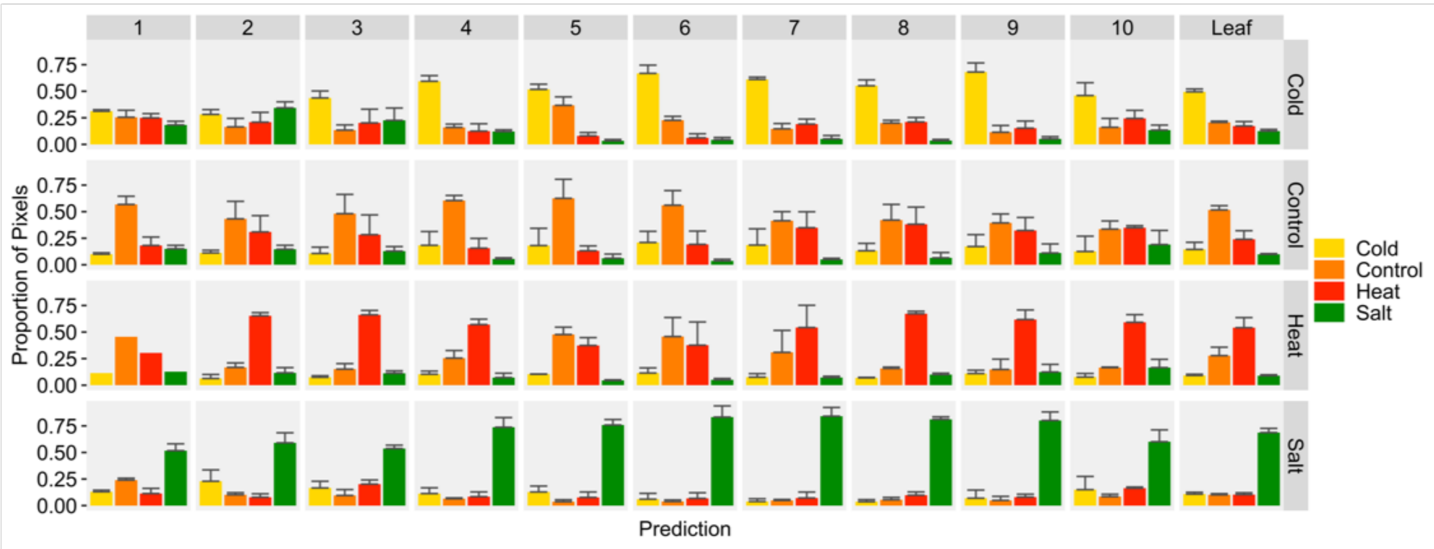


**Figure S6.** Average proportion of pixels from all MS71 (**A**) and PH207 (**B**) plants of a certain treatment (rows) for each leaf segment (columns) predicted to belong to a certain treatment class. Bars represent the mean proportion of all plants per category and error bars represent the standard deviation around the mean.

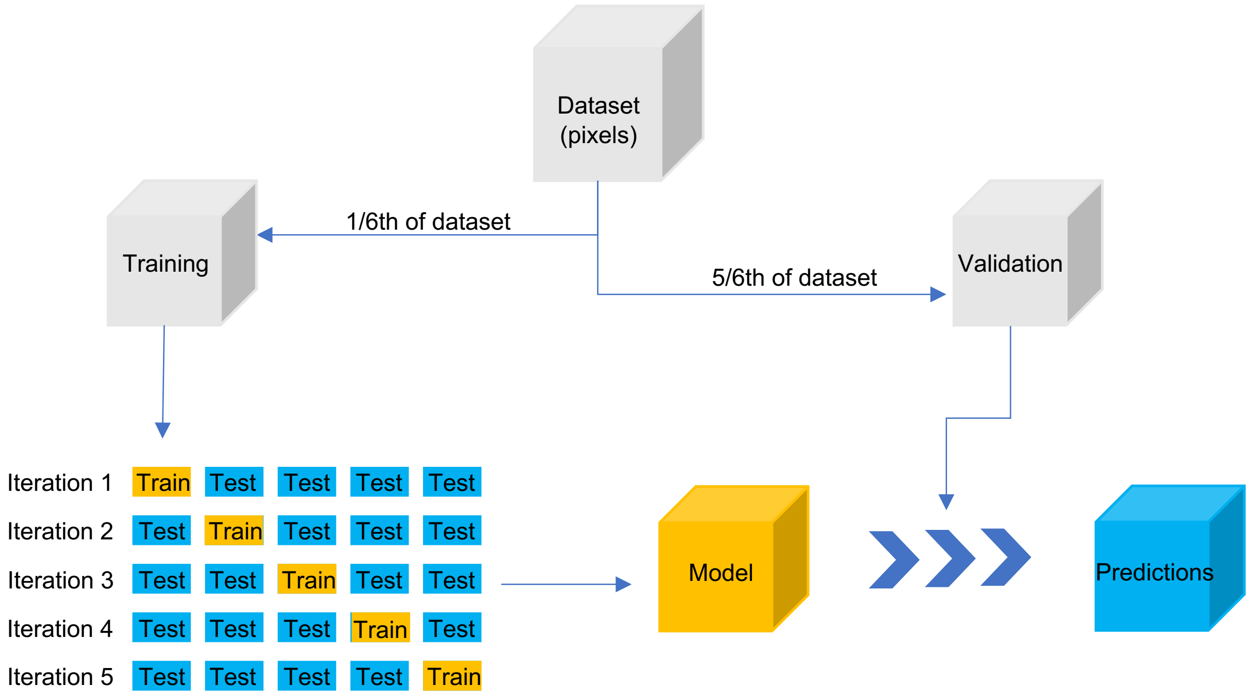


**Figure S7. SVM model training and testing procedure.**
